# Host-specific microbiome and genomic signatures in *Bifidobacterium* reveal co-evolutionary and functional adaptations across diverse animal hosts

**DOI:** 10.1101/2025.01.10.632322

**Authors:** Magdalena Kujawska, David Seki, Lisa Chalklen, Jennifer Malsom, Sara Goatcher, Ioulios Christoforou, Suparna Mitra, Lucy Crouch, Lindsay J Hall

## Abstract

Animal hosts harbour divergent microbiota, including various *Bifidobacterium* species and strains, yet their evolutionary relationships, and functional adaptions remain understudied. By integrating taxonomic, genomic and predicted functional annotations, we uncover how *Bifidobacterium* adapts to host-specific environments, shaped by vertical transmission, dietary influences, and host phylogeny. Our findings reveal that host phylogeny is a major determinant of gut microbiota composition. Distinct microbial networks in mammalian and avian hosts reflect evolutionary adaptations to dietary niches, such as carnivory, and ecological pressures. At a strain-resolved level, we identify strong co-phylogenetic associations between *Bifidobacterium* strains and their hosts, driven by vertical transmission and dietary selection, underscoring the intricate co-evolutionary dynamics between these microbes and their hosts. Functional analyses highlight striking host-specific metabolic adaptations in *Bifidobacterium*, particularly in carbohydrate metabolism and oxidative stress responses. In mammals, we observe an enrichment of glycoside hydrolases (GH) tailored to complex carbohydrate-rich diets, including multi-domain GH13_28 α-amylases featuring diverse carbohydrate-binding modules (CBM25, CBM26, and the novel CBM74). These adaptations emphasise the ecological flexibility of *Bifidobacterium* in breaking down α-linked glucose polysaccharides, such as resistant starch. Together, our study provides new insights into the evolutionary trajectories and ecological plasticity of *Bifidobacterium*, revealing how host phylogeny and dietary ecology drive microbial diversity and function. These findings deepen our understanding of host-microbe co-evolution and the critical role of microbiota in shaping animal health and adaptation.

## Introduction

Members of the genus *Bifidobacterium* have long been associated with beneficial health outcomes, particularly due to their probiotic properties. Human-derived bifidobacteria, especially those associated with infancy, have received considerable research attention, leading to extensive efforts in their isolation and characterisation (1–3). However, *Bifidobacterium* species are not confined to humans; they are widely distributed across the animal kingdom, associating with a diverse array of hosts belonging to different classes, including mammals, birds, and social insects (4,5). Increased availability and affordability of next-generation sequencing (NGS) have significantly expanded our understanding of *Bifidobacterium* diversity, revealing novel species and elucidating inter- and intra- species variability.

The distribution of *Bifidobacterium* across diverse hosts has led to the evolutionary hypotheses regarding their host specificity. Some species like *Bifidobacterium* animalis and *Bifidobacterium* pseudolongum, appear to be generalist, found across diverse animal classes. In contrast, other species demonstrate stronger host specificity, particularly within primates (6,7). However, research has been disproportionately dominated by observations from primates, with approximately half of the recognised *Bifidobacterium* species type strains recovered from non-human primates, particularly members of the family Callitrichidae (8,9). This created a bias in our understanding of bifidobacterial host range and ecological specialisation, even though other mammalian hosts, and indeed other classes of animals, likely harbour a wealth of undiscovered *Bifidobacterium* species and subspecies (10–12).

Central to evolutionary studies is the concept of phylosymbiosis, which refers to the evolutionary concordance between host phylogeny and microbiome composition (13). This phenomenon suggests that closely related host species tend to harbour more similar microbial communities, likely due to co-evolutionary processes. The existence of such processes between two, or more, taxa can be assessed by co-phylogeny, which evaluates the congruence of phylogenetic relationships between different groups of organisms due to their long-term interaction (14–16). Despite the growing availability of genomic data, robust studies investigating evolutionary and functional relationships between *Bifidobacterium* and their animal hosts at the strain level remain scarce. The pan-genome of the genus and its phylogenetic relationship are yet to be fully resolved. Co-phylogenetic analyses have suggested a close evolutionary relationship between certain bifidobacteria and their primate hosts, particularly those within the Hominidae family (17,18). Previous large-scale studies have often focused on intra-species features or a limited number of isolates, leaving much to be explored about the genomic and functional diversity of *Bifidobacterium* across the animal kingdom (19,20).

In the context of *Bifidobacterium*, phylosymbiosis could play a crucial role in shaping the distribution and functionality of these microbes across different hosts. Additionally, diet plays a critical role in the shaping of the microbiome, with glycoside hydrolases (GH) – enzymes which break down complex carbohydrates – particularly important (21,22). The diversity and evolution of GH enzymes within *Bifidobacterium* genomes offer insights into how diet and microbial function co-evolve in a host- specific manner, driving adaptation and ecological flexibility (23).

To address existing knowledge gaps and expand our understanding of *Bifidobacterium* diversity, we sequenced a total of 230 faecal samples from a broad range of animal hosts, including representatives from the classes Aves, Reptilia, Insecta, and Mammalia. In addition, we isolated and sequenced 96 unduplicated *Bifidobacterium* strains, which we used as basis to compile a robust genomic dataset supplemented by high-quality public data. We conducted a comprehensive analysis of the host gut microbiota composition and *Bifidobacterium* diversity and functions, particularly those related to carbohydrate metabolism and the most abundant GH13 family, to assess the evolutionary links between diet, microbial function, and host phylogeny. This study aims to provide a more detailed understanding of the adaptive strategies that enable *Bifidobacterium* to thrive across diverse host environments, offering new insights into the co-evolution of hosts and their microbiomes.

## Materials and methods

### Sample collection

Animal faecal samples were collected by animal-care staff at Banham Zoo (UK), Africa Alive! (UK) and Pafos Zoo (Cyprus) into sterile Sterilin (Thermo Scientific) specimen containers with a spoon. Samples from Banham Zoo and Africa Alive! were stored at 4°C under anaerobic conditions using Oxoid AnaeroGen 2.5L sachets and transported to our laboratory within 48h. Samples from Pafos Zoo were kept in -20°C until shipped to the UK. At our laboratory, all samples were stored at -80°C.

### Isolation of bifidobacteria

Depending on the amount of available faecal material, samples (∼50mg or ∼100mg) were re- suspended in either 450μl or 900μl of sterile Phosphate Buffer Saline, vortexed for 30s, mixed on a shaker at 1600rpm, and used to produce serial dilutions (neat to 10^-4^). The dilutions were plated onto either de Man-Rogosa-Sharpe (MRS) agar (BD Biosciences) supplemented with mupirocin (50mg/l) (AppliChem) and L-cysteine hydrochloride monohydrate (50mg/l) (Sigma-Aldrich) or Brain Heart Infusion (BHI) agar (Oxoid) supplemented with mupirocin (50mg/l), L-cysteine hydrochloride monohydrate (50mg/l) and sodium iodoacetate (7.5mg/l) (Oxoid). Plates were incubated in an anaerobic cabinet (Don Whitley Scientific) for 48-72 hours. Three colonies from each dilution were randomly selected and streaked to purity on either BHI agar or MRS agar supplemented with L- cysteine hydrochloride monohydrate (50mg/l). Pure cultures were stored in cryogenic tubes at -80°C.

### DNA extraction from faecal samples

FastDNA Spin Kit for Soil (MP Biomedicals) was used to extract DNA from ∼200mg of animal faecal material following manufacturer instructions, with the bead-beating time extended to 3 min. DNA concentration and quality were quantified using Qubit dsDNA BR Assay Kit in Qubit 2.0 Fluorometer according to the manufacturer’s protocol (Invitrogen).

### 16S rRNA amplicon sequencing and data analysis

We sequenced a total of 230 animal faecal samples using the V1-V2 16S rRNA gene region (n= 21) and the V4 16S rRNA gene region (n = 209). PCR amplicons for the V1-V2 region of the 16S rRNA gene were generated with primers: Modified 27F 5’- AGMGTTYGATYMTGGCTCAG-3’ (24) and 338R 5’- GCTGCCTCCCGTAGGAGT-3’ (25), and sequenced at Novogene (Cambridge, UK). PCR amplicons for the V4 region were generated with primers SA501-SA508(F) and SA701-SA712(R) (26) and sequenced at Earlham Institute (Norwich, UK). Cutadapt v.1.18 (27) was used to remove primers from each dataset separately. Reads below the quality score of 20 (minQ=20) and maximum expected error of 1 (maxEE=1) were filtered out from each dataset using DADA2 (28), followed by the inference of amplicon sequence variants (ASV). Chimeras were removed from each dataset separately, after which the datasets were merged. Taxonomy was assigned using SILVA database v.138 (29). ‘Ampvis2’ (30) package implemented in R v.4.2.3 (31) was used to rarefy the ASV table at 10,000 reads per sample and to analyse and visualise the 16S rRNA amplicon data from hosts belonging to classes Insecta, Reptilia, Aves and Mammalia (n = 219 samples, Supplementary Table 1). Data from Actinopterygii and Gastropoda were not included in the analysis. PERMANOVA tests were performed using the R package ‘vegan’ (32). Hierarchical clustering was performed with ‘pheatmap’ (33). All p-values were adjusted using Bonferroni’s method. Data was visualized using the R package ‘ggplot2’ (34).

To perform phylosymbiosis analysis using the 16S rRNA amplicon data, we first used the TimeTree v.5.0 resource (35) to acquire separate dated phylogenetic trees encompassing as many mammalian and avian host species from out dataset as possible (62 mammalian and 32 avian hosts). We then obtained the microbiota composition of each host by averaging the corresponding sample 16S rRNA amplicon data per host species and extracted the relative abundances of the main bacterial orders. We only took the 15 most abundant orders under consideration. We used the ‘ABDOMEN’ (36) R suite to apply the model of microbiota evolution separately to mammalian and avian datasets, with 3000 STAN (37) iterations permutated 10 times.

### Genomic DNA extraction from bacterial isolates

Bacterial pellets were re-suspended in 2ml of 25% sucrose in 10mM Tris (Sigma-Aldrich) and 1mM EDTA (Sigma-Aldrich) at pH8.0. Cells were then treated using 50μl of 100mg/ml lysozyme (Roche). Further, 100μl of 20mg/ml Proteinase K (Roche), 30μl of 10mg/ml RNase A (Roche), 400μl of 0.5lllM EDTA and 250μl of 10% Sarkosyl NL30 (Sigma-Aldrich) were added into the lysed bacterial suspension. The samples were then incubated on ice for 2 hours, followed by 50°C overnight water bath.

Next, samples were subject to three rounds of Phenol:Chloroform:Isoamyl Alcohol (25:24:1) (Sigma-Aldrich) extraction using Qiagen MaXtract High Density tubes. Further two rounds of extractions with Chloroform:Isoamyl Alcohol (24:1) (Sigma-Aldrich) were then performed to remove residual phenol, followed by ethanol precipitation and 70% ethanol wash, after which DNA pellets were resuspended in 300μl of 10mM Tris (pH8.0). Sample DNA concentration was quantified using Qubit dsDNA BR Assay Kit in Qubit 2.0 Fluorometer. Extracted DNA was stored in -20°C until further analysis.

### Whole genome sequencing

This work was performed at the Wellcome Trust Sanger Institute (Hinxton, UK) and at the Quadram Institute Bioscience (Norwich, UK). At Hinxton, DNA was sequenced using 96-plex Illumina HiSeq 2500 platform as described previously (3). At Quadram Institute Bioscience, genomic DNA was normalised to 0.5ng/µl with EB (10mM Tris-HCl). 0.9µl of TD Tagment DNA Buffer (Illumina Catalogue No. 15027866) was mixed with 0.09µl TDE1, Tagment DNA Enzyme (Illumina Catalogue No. 15027865) and 2.01µl PCR grade water in a master mix and 3μl added to a chilled 96 well plate. 2µl of normalised DNA (1ng total) was pipette mixed with the 3µl of the tagmentation mix and heated to 55 ⁰C for 10 minutes in a PCR block. A PCR master mix was made up using 4μl kapa2G buffer, 0.4µl dNTPs, 0.08µl Polymerase and 6.52µl PCR grade water, contained in the Kap2G Robust PCR kit (Sigma Catalogue No. KK5005) per sample and 11µl added to each well need to be used in a 96-well plate. 2µl of each P7 and P5 of Nextera XT Index Kit v2 index primers (Illumina Catalogue No. FC-131-2001 to 2004) were added to each well. Finally, the 5µl of tagmentation mix was added and mixed. The PCR was run with 72⁰C for 3 minutes, 95⁰C for 1 minute, 14 cycles of 95⁰C for 10s, 55⁰C for 20s and 72⁰C for 3 minutes. Following the PCR reaction, the libraries were quantified using the Quant-iT dsDNA Assay Kit, high sensitivity kit (Catalogue No. 10164582) and run on a FLUOstar Optima plate reader. Libraries were pooled following quantification in equal quantities. The final pool was double-SPRI size selected between 0.5 and 0.7X bead volumes using KAPA Pure Beads (Roche Catalogue No. 07983298001). The final pool was quantified on a Qubit 3.0 instrument and run on a High Sensitivity D1000 ScreenTape (Agilent Catalogue No. 5067-5579) using the Agilent Tapestation 4200 to calculate the final library pool molarity. The pool was run at a final concentration of 10pM on an Illumina MiSeq instrument.

### Genomic data processing

For data generated at the Wellcome Trust Sanger Institute, genome assemblies were performed by the sequencing provider using the assembly pipeline described by Page et al. (38). Sequencing reads generated at the Quadram Institute Bioscience were pre-processed with fastp v.0.23.2 (39) and assembled using Unicycler v.0.4.9 (40) with the “--mode conservative” option, after which contigs below 1000lllbp were filtered out. We estimated completeness and contamination of all genomes sequenced as part of this study using CheckM v.1.2.0 (41), and retained sequences with completeness >99% and contamination <1%.

Additionally, genomes of animal-associated *Bifidobacterium* isolates were identified and downloaded from the NCBI. dRep v.2.5.0 was used to dereplicate the dataset at 99.9% identity threshold (42).

GTDB-Tk v.2.1.0 (43) was used to classify all genomic sequences to the strain level and produce a multiple sequence alignment of the single copy gene markers. Python3 module pyANI v.0.2.10 with default settings was used to calculate the average nucleotide identity values (ANI) (44). Species delineation cut-off was set at 95% identity (45). After such pre-processing, we retained 387 *Bifidobacterium* genomic sequences as a final dataset.

### Phylogenetic analysis

IQ-Tree v.2.0.5 (46) was used to test for the best substitution model fitting the GTDB-Tk-created alignment and produce a global *Bifidobacterium* phylogenetic tree, which was visualised with ITOL v.6.9 (47).

### Co-phylogeny analysis

TimeTree v.5.0 resource was used to retrieve dated trees for mammalian and avian hosts harbouring bifidobacteria from our dataset. GTDB-Tk and IQ-Tree were then used to produce alignments and phylogenies for bifidobacteria associated with these mammalian and avian hosts, respectively. ‘Parafit’ function from the ‘ape’ R package (48) with the p-value significance level set at 0.01 and the permutation number set to 999 was run 3 times to test for the host-*Bifidobacterium* co-phylogenetic signal. P-values obtained from each run were adjusted using the Benjamini-Hochberg correction. Tanglegrams were created with ‘phytools’ (49) package implemented in R.

### Genomic and functional annotation

For consistency, all genomes were annotated using Prokka v.1.14.6 (50). Functional traits were profiled using eggNOG-mapper v.2.1.11 (51) with eggNOG database v.5.0 (52). Standalone version of dbCAN3 with HMMdb v.12 and “--hmm_cov 0.50” option was used to annotate carbohydrate-active enzymes (CAZYmes). R packages ‘dplyr’ (53) and ‘ggplot2’ were used to summarise and visualise basic genomic and functional abundance features.

### Assessment of between-group differences in abundance of functional features

Between-group analysis of differences in abundance of functional features in bifidobacteria associated with particular host groups was performed in R for a subset of the genomic data. We only included host order categories for which we had more than 20 *Bifidobacterium* genomes available (Hymenoptera (n = 35), Artiodactyla (n = 58), Carnivora (n = 22), Rodentia (n = 47), Primates (n = 172)). KO and CAZyme abundance matrices were filtered at mean prevalence of >20% and subsequently log-transformed (log+1) to achieve a less skewed distribution. Differences were assessed following the approach described in Ruehlemann et al. (54). Briefly, linear regression analysis was employed using abundances as dependent variable and Hymenoptera/Mammals and Primate/non-primate dichotomies as explanatory variables in a single model for each function defined as lm(abundance ∼ Mammals + Primates). P-values were calculated from the t-values of the resulting models using the summary.lm() function. Log-fold differences were calculated using group mean abundances and a pseudo count of 0.01. P-values were adjusted using Bonferroni correction. Features with significant (Qlll<lll0.05) positive association were grouped into “Hymenoptera”, “Mammals”, “Primates” and “non-primates” categories. Features without abundance differences were grouped as “other”.

Additionally, CAZyme abundance data for host order groups was summarised and visualised with R packages ‘dplyr’, ‘ggplot2’, ‘cowplot’ (55) and ‘UpSetR’ (56). Principal component analysis of the CAZyme abundance data was performed using ‘factoextra’ v.1.0.7 (57).

### Prediction of protein structure and ligand binding of selected glycoside hydrolases

AlphaFold 3 available through the Google DeepMind platform was used to predict protein structures of selected glycoside hydrolases based on their amino acid sequences. Reference protein models coupled with ligands were downloaded from the Protein Data Bank (58). We used Coot v.0.9.8.93 to align models produced in this study to reference models (59). PyMOL v.3.0.0 was used to visualise superimposed structures (60). PeSTo-Carbs was used to predict carbohydrate binding sites in protein structures, to which we were not able to align available reference ligands (61).

## Results

### Host-specific microbial diversity and enrichment of *Bifidobacterium* in animal gut microbiomes

Here, we investigated the taxonomic composition of the gut microbiota across diverse animal classes, to identify patterns of microbial diversity and the specific association of *Bifidobacterium* with certain host lineages. We analysed a total of 219 faecal samples from 126 diverse animal hosts, including representatives of classes Insecta, Reptilia, Aves and Mammalia, using 16S rRNA amplicon sequencing data.

After rarefying the dataset to 10,000 reads per sample and filtering out low-abundance amplicon sequencing variants (ASVs) occurring at < 1%, we conducted a principal component analysis to assess differences in bacterial diversity between host orders (Figure 1A). Overall, analysis revealed significant differences in microbiota composition according to host order (PERMANOVA, p.adj < 0.05), with beta diversity significantly different between the taxonomic classes Mammalia and Aves, Reptilia, and Insecta (PERMANOVA, p.adj = 0.006, 0.04, and 0.04 respectively). Pairwise comparisons of microbial composition between host orders highlighted that taxa associated with Artiodactyla were significantly different from those found in other lineages (Figure 1B), forming a distinct cluster as seen in Figure 1A. Furthermore, microbial richness varied greatly across different host orders, with the lowest richness observed in Bucerotiformes, and the highest in Marsupialia (Figure 1C). Notably, Escherichia coli was prevalent and dominant in the majority of avian host orders, except for Coraciiformes and Struthioniformes. *Bifidobacterium* ASVs were particularly enriched in Edentata and Primates (Figure 1D). This observation underscores the potential evolutionary and ecological significance of *Bifidobacterium* in these groups. Further analysis using hierarchical clustering of bacterial genera by host order showed that carnivores cluster closely to birds of prey (Falconiformes and Accipitriformes). In contrast, primates displayed a distinct association with *Bifidobacterium* (Fig. 1E), hosting the highest relative abundance of ASVs corresponding to this genus. The abundance of *Bifidobacterium* in primates was significantly higher than in the gut microbiomes of Squamata, Artiodactyla, and Carnivora (ANOVA, p = 0.01, 0.001, and <0.0001, respectively) (Fig. 1E). This pronounced association appears to be driven by very high abundance of bifidobacteria in tamarins and marmosets (Figure 1F), which also exhibit notably low alpha diversity compared to other New World monkeys (Figure 1G). These findings suggest that certain primate species, particularly those within the Callitrichidae family, may have evolved specialised gut environments that support *Bifidobacterium* persistence, possibly linked to their diet and unique ecological niches.

**Fig. 1.**
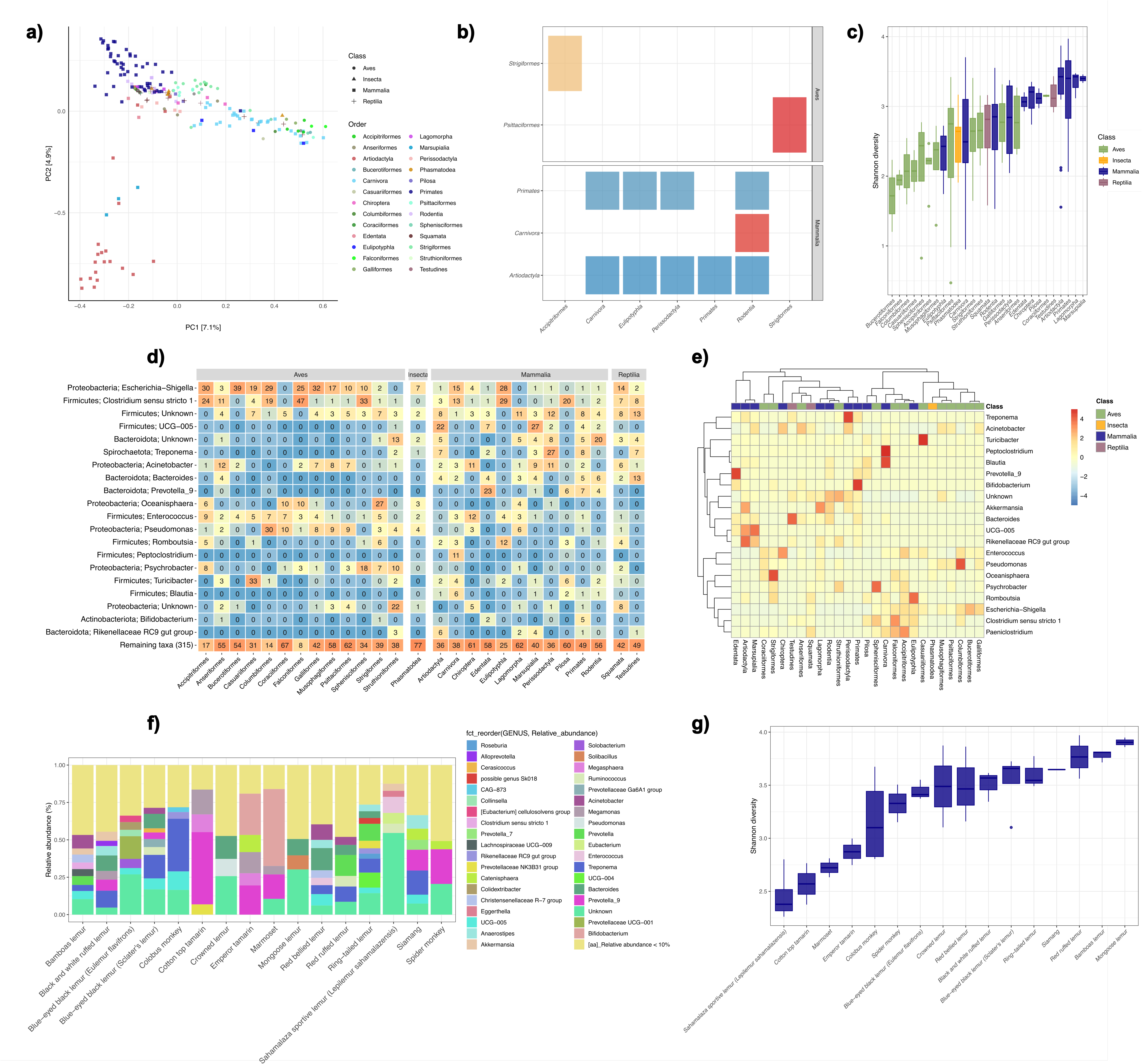
Characteristics of gut microbiota in insects, reptiles, birds, and mammals. **a)** Principal component analysis (PCA) of relative species counts. **b)** Heatmap displaying significant results (PERMANOVA, p.adj < 0.05) out of all pairwise comparisons. Blue colour indicates low Euclidean distance, red colour indicates high Euclidean distance between given host pairs. **c)** Microbial richness measured by Shannon’s index. **d)** Heatmap displaying mean average abundance of species in given hosts. Blue colour indicates low mean average abundance, orange colour indicates high mean average abundance. **e)** Hierarchical clustering of pairwise complete observations between host and occurrence of microbial species using Pearson correlation. **f)** Stacked bar plot displaying relative abundance of microbial genera in primates. **g)** Microbial richness in primate hosts measured by Shannon’s index.

### Phylosymbiosis signals and microbial covariance patterns across mammalian and avian lineages

Microbiota composition can vary significantly within and between host species (62,63), yet more closely related hosts often harbour a more similar microbiota (13,64), a phenomenon known as phylosymbiosis. To investigate phylosymbiosis patterns in our dataset, we applied a multivariate Brownian motion model (36) to the gut bacterial microbiota of 62 mammalian species and 38 bird species. We focused on the 15 most abundant bacterial orders to evaluate the strength of the phylogenetic signal in these groups. Our analysis revealed a stronger phylosymbiosis signal in mammals compared to birds (λ ≃ 0.41 and λ ≃ 0.24, respectively). This suggests that the gut microbiota of mammals is more tightly linked to host phylogeny than that of birds. Ancestral reconstruction indicated that members of phylum Pseudomonadota, particularly the order and Enterobacteriales, were more abundant in the ancestral gut microbiota of birds than mammals (Fig.2a and b). This distinction likely reflects differences in the evolutionary history and dietary habits of these two host groups. This analysis also revealed notable shifts in bacterial community composition in the ancestors of various mammalian and avian orders. In mammals, the largest shift was detected in the ancestor of Carnivora, characterised by an increased proportion of Clostridiales (Bacillota), Peptostreptococcales – Tissierellales (Bacillota) and Enterobacteriales (Pseudomonadota), along with a decreased proportion of Bacteroidales (Bacteroidota). Another notable shift occurred in the ancestor of the Primate suborder Simiiformes (encompassing both New and Old World monkeys), with exhibited an increased proportion of Bacteroidales. Additionally, an increased proportion of Veillonellales – Selenomonadales was detected in the ancestor of New World monkeys. In birds, a major shift was observed in the ancestor of clade Neoaves, marked by an increased proportion of Enterobacteriales (Pseudomonadota). In addition, the ancestors of Falconiformes and Sphenisciformes were characterised by a larger proportion of Clostridiales (Bacillota), highlighting a possible adaptation to their carnivorous diets and unique ecological niches.

Consistent with previous reports, we detected both positive and negative covariances among various bacterial taxa in mammals and birds. Visual inspection of the covariance matrices revealed the clustering of bacterial orders into subsets, with taxa within these subsets tending to covary interdependently. In both mammals and birds, positive covariances were observed between bacterial members of Pseudomonadota and Bacilllota, with this covariance profile particularly dominant in the microbiota of Carnivora and carnivorous birds (Fig. 2a and b). In mammals, notably, Bacteroidales and Coriobacteriales covaried with Bifidobacteriales and Veillonellales-Selenomonadales in New World monkeys alone, suggesting a unique microbiota profile in these primates (Fig. 2a). In birds, we detected strong covariance between members of Actinomycetota and Pseudomonadota, with this subset exhibiting a strong negative covariance with Clostridiales (Fig. 2b). These results indicate the existence of distinct microbial networks in both mammalian and avian hosts that may reflect their evolutionary adaptations, possibly driven by host diet and collaborative nature of microbial interactions.

**Fig. 2.**
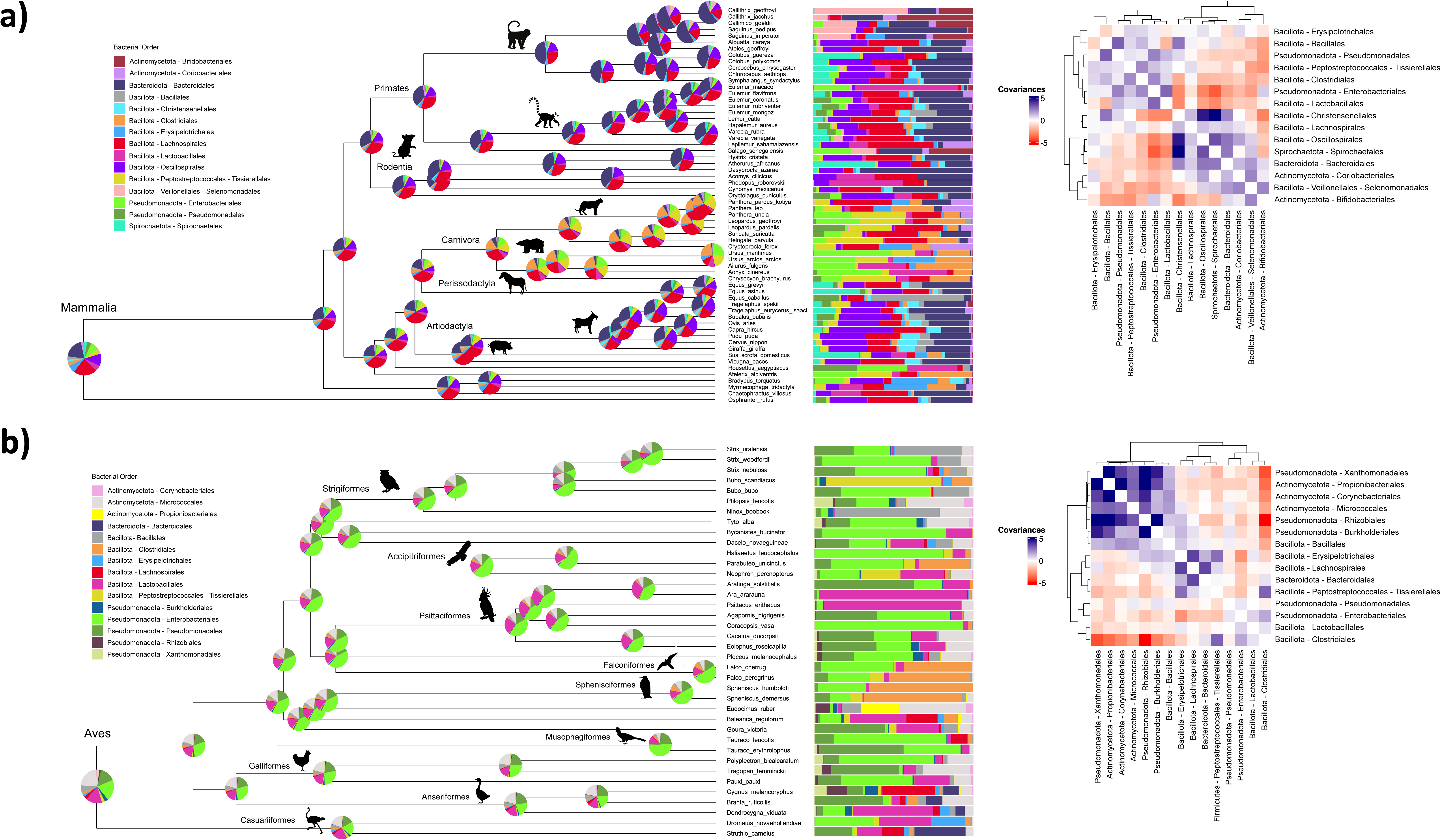
Ancestral reconstruction of mammalian **a)** and avian **b)** gut microbiota with estimated covariances between the main bacterial taxa that tend to be similar in the gut microbiota of mammals and birds (heatmaps on the right). Phylogenetic tree of the sampled mammalian and avian hosts and associated relative abundances of the 15 most abundant bacterial orders (bar charts to the right of phylogenies) are shown. Pie charts at the root and nodes of the tree represent estimated ancestral microbiota compositions. For each covariance matrix, we represented negative covariances in red and positive covariances in blue.

### Genomic diversity, phylogenetic congruence, and general functional features of animal-associated *Bifidobacterium*

To expand current understanding of genomic features of animal-associated bifidobacteria, we isolated and sequenced strains from a broad range of hosts across the animal tree of life (Supplementary Table 2). We successfully recovered *Bifidobacterium* from 37 out of 175 (21.1%) host species included in the study, with the highest isolate recovery rate observed in mammals (27 out of 87 hosts, 31%), followed by birds (8 out of 59 hosts, 13.6%) and reptiles (2 out of 15 hosts, 13.3%). Our efforts resulted in the final collection of 96 unduplicated bifidobacterial strains from mammals (n=84), birds (n=10) and reptiles (n=2). The genome sizes of these isolates ranged from 1.85 Mb (*Bifidobacterium* pseudolongum subsp. pseudolongum) to 3.26 Mb (*Bifidobacterium* simiventris), corresponding to 1,529 and 2,674 protein-coding open reading frames (ORFs), respectively. These values fall within the range for previously reported for *Bifidobacterium* species (4,23).

Using GTDB-tk analysis and comparison of ANI values between the recovered 96 isolates and 105 *Bifidobacterium* type strains, we assigned 66 genomes (69%) to 32 *Bifidobacterium* species and subspecies (ANI > 95%) (Supplementary Table 3 and Supplementary Table 4). The predominant species and subspecies identified included *Bifidobacterium* pseudolongum subsp. globosum (7 isolates, 11% of assigned genomes), *Bifidobacterium* reuteri (6 isolates, 10% of assigned genomes) and *Bifidobacterium* animalis subsp. animalis (4 isolates, 6% of assigned genomes) and *Bifidobacterium* callitrichidarum (4 isolates, 6% of assigned genomes). Surprisingly, 30 isolates (31%) showed ANI values below 95% compared to known *Bifidobacterium* type strains, suggesting they may represent putative novel species. These novel isolates were primarily recovered from primates (63%), with other hosts including a South American rodent – Dasyprocta azarae (Azara’s agouti), various bird species, and a red-footed tortoise (Chelonoidis carbonaria). ANI comparisons between these novel isolates indicated that they belonged to 11 putative novel species.

To further explore the diversity within the *Bifidobacterium* genus, we expanded our dataset by including 291 publicly available animal-associated genomes, resulting in a total dataset of 387 strains (Fig.3). Phylogenetic analysis revealed moderate clustering of *Bifidobacterium* strains according to host order, supported by ANOSIM statistics (R = 0.5037, p = 0.0001). Strains isolated from the hosts belonging to same order tended to cluster together within bifidobacterial species (Fig.3a), reflecting the notion that certain species such as *Bifidobacterium* asteroides, are host specific (e.g. only found in insects), while others like B. animalis or B. pseudolongum, exhibit more cosmopolitan distributions. We further tested for co-phylogeny signal between mammalian and avian host phylogenies and those of their associated bifidobacterial strains, using a global-fit approach implemented in the R function ‘parafit’. Significant phylogenetic congruence was detected in Mammalia (ParaFitGlobal = 10071739, mean p_global_ = 0.001, but not in Aves (ParaFitGlobal = 21600.47, mean p_global_ = 0.122) (Fig. 3b). Most mammals displayed significant links to associated strains (Supplementary Table 5). Notable exceptions included the African elephant, sloth, domestic cat, fruit bat and certain primate species. Interestingly, for the order Artiodactyla, phylogenetic congruence with associated strains seemed to be dependent on the specific *Bifidobacterium* species, with significant associations found between “cosmopolitan” species like B. pseudolongum and their porcine host.

**Fig. 3.**
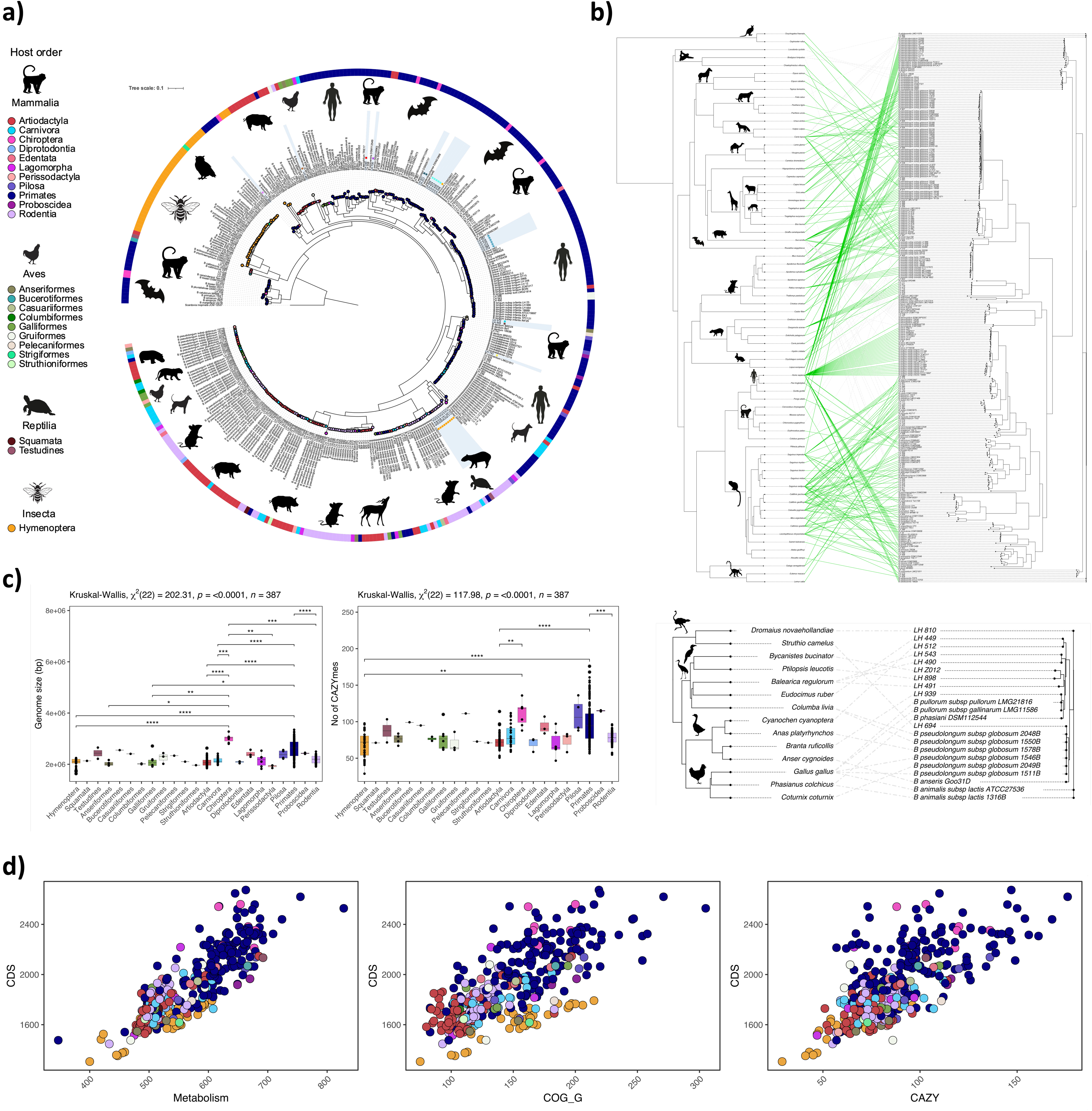
General features of bifidobacterial genomes used in this study, co-phylogeny analysis, and general functional traits of animal-associated bifidobacteria. **a)** Maximum likelihood tree of 387 animal-associated bifidobacterial genomes. Isolation source (animal host order) is marked with a coloured strip and node symbols on the tree. Light blue shading marks strains belonging to potential novel species, and coloured squares in the shading denote individual novel species. **b)** Tanglegram reflecting co-phylogeny of bifidobacterial strains and their hosts. Significant links are marked in green. **c)** Comparison of genome sizes and carbohydrate-active enzyme (CAZyme) abundance between **Bifidobacterium** isolates grouped according to host order. Significant differences between host-order-associated groups based on the Kruskal-Wallis statistics are denoted with asterisks. **d)** Number of coding sequences as a function of number of genes involved in overall metabolism (left), carbohydrate metabolism (middle) and representing CAZyme abundance in bifidobacterial genomes grouped according to host order.

Comparative analysis of genome sizes across *Bifidobacterium* isolates grouped by host order indicated significant differences (Kruskal-Wallis x^2^ = 202.31, P < 0.0001, df = 22). For example, *Bifidobacterium* isolates (Fig.3c) from Chiroptera (bats) and Primates had the largest genomes (3.01 ± 0.16 Mb and 2.60 ± 0.33 Mb (mean ± sd), respectively), while those associated with Perissodactyla (horses, rhinoceroses) and Struthioniformes (ostriches) had the smallest genomes (1.92 ± 0.1 Mb and 1.96 ± 0 Mb, respectively). These findings align with previous analyses of 129 publicly available *Bifidobacterium* strains (20).

Functional classification of gene content assigned 80.9% of ORFs to Clusters of Orthologous Groups (COG) categories, with 19.1% categorised as proteins of unknown function (Supplementary Table 6). As expected for *Bifidobacterium*, carbohydrate transport and metabolism (COG_G) was the second most abundant category (after function unknown) and constituted 9.5% of assigned functions, reflecting the saccharolytic lifestyle of this genus. Additionally, genes involved in amino acid metabolism (COG_E) constituted 8.6% of assigned functions. Carbohydrate active enzymes (CAZymes) made up 59.1% of ORFs in the COG_G category, with glycoside hydrolases being the predominant class across all host orders. Notably, the abundance of putative GH enzymes varied significantly between host orders (Kruskal-Wallis x^2^ = 117.98, P < 0.0001, df = 22), with a tendency for strains with larger genomes to harbour more carbohydrate metabolism genes (Fig. 3c and d).

### Host-specific carbohydrate metabolism adaptations in *Bifidobacterium*

Given that taxon-specific functional properties can reflect broad-scale differences between host groups, we conducted a focused analysis on the functions of *Bifidobacterium*, particularly those related to carbohydrate metabolism. We concentrated on host order groups for which we had more than 20 *Bifidobacterium* genomes available; Hymenoptera, Artiodactyla, Carnivora, Rodentia, Primates.

We analysed 1,555 KEGG orthologues (65) to identify abundance differences between *Bifidobacterium* from Hymenoptera vs. those associated with mammals, both Primates and non-Primates. We found significant abundance differences in 377 (24.2%) and 292 (18.7 %) KOs, respectively (QBonferroni<0.05; Fig. 4a, Supplementary Table 7). Specifically, bifidobacterial genomes associated with insect hosts (Hymenoptera) exhibited higher abundances of KOs related to oxidative phosphorylation, particularly components of the cytochrome bd system. In contrast, mammal-associated *Bifidobacterium* showed higher abundance in categories linked to ferrous iron transport, cofactor and vitamin biosynthesis and carbohydrate metabolism, notably including enzymes belonging to GH13_9 subfamily (K00700), where the current members have been characterised as hydrolysing the α-1,6-branches in amylopectin and glycogen. Primate-associated genomes were enriched in categories covering major facilitator superfamily transporters, glycerophospholipid metabolism and metabolism of cofactors and vitamins.

**Fig 4.**
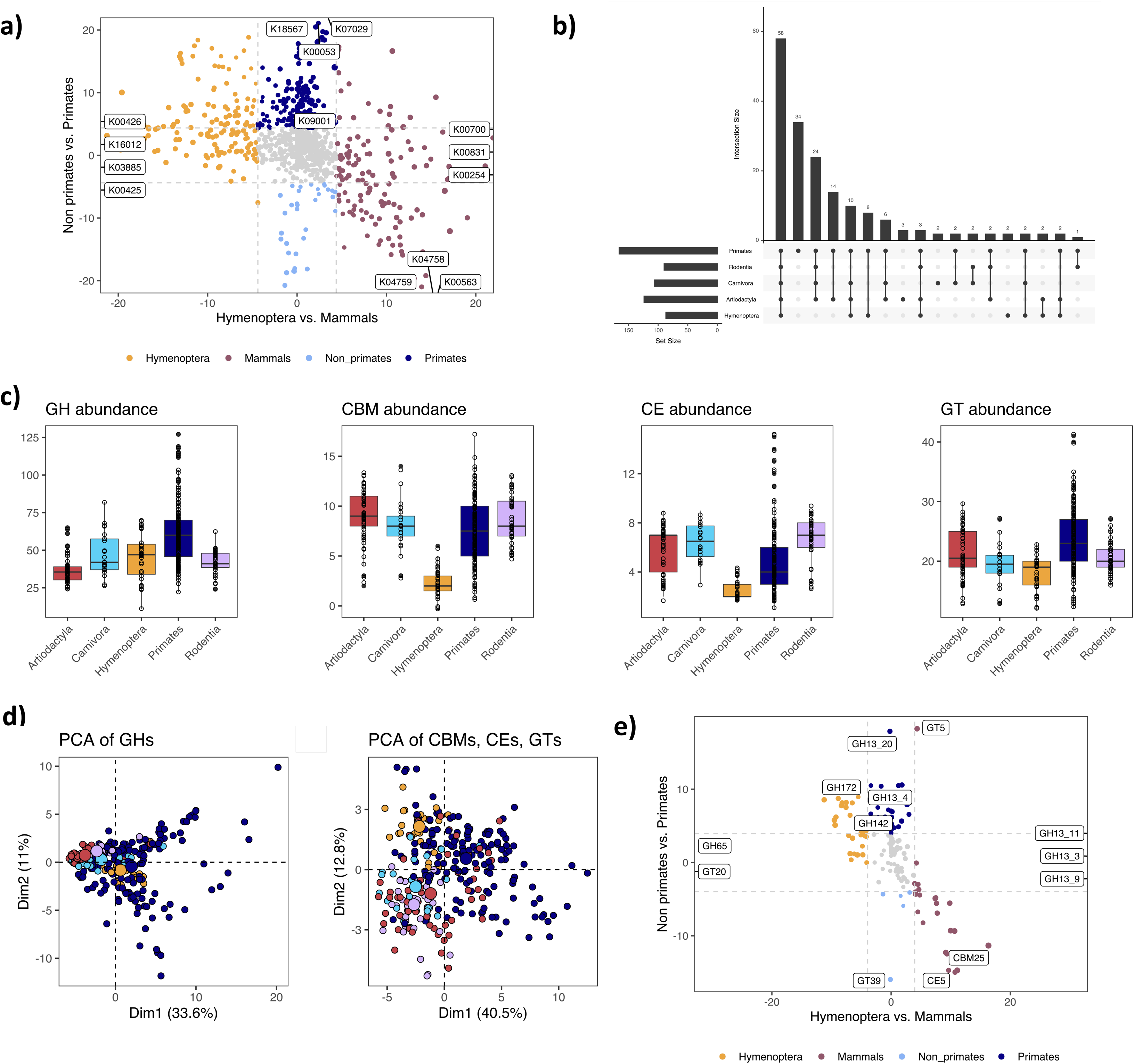
Functional analysis of bifidobacterial genomes from the perspective of host taxonomic level of order. **a)** Analysis of abundance differences of 1,555 KEGG orthologs in **Bifidobacterium** from Hymenoptera vs. those associated with mammals, Primates and non-primates. Top KO categories associated with each group are marked on the plot. **b)** Distribution of CAZyme classes in bifidobacteria grouped according to host order groups. **c)** Abundance of CAZyme classes in bifidobacterial genomes grouped according to host order; GH – glycoside hydrolase, CBM – carbohydrate binding module, CE – carbohydrate esterase, GT – glycosyltransferase. **d)** PCA analysis based on the CAZyme abundance matrix, with data for glycoside hydrolases analysed separately. **e)** Analysis of abundance differences of 189 CAZyme classes in **Bifidobacterium** from Hymenoptera vs. those from mammals, Primates, and non-primates. Top CAZyme classes associated with each group are marked on the plot.

We further examined the distribution of CAZyme families and subfamilies across host order groups. It is important to note that assigning a putative enzyme to a glycoside hydrolase (GH) family does not always provide a definitive indication of its activity, as this can be highly family-dependent. Currently, there are 189 GH families, with substantial variability in the number of characterised members per family. Some families have hundreds of well-studied enzymes, while others may only have a single characterised member. For example, enzymes assigned to the relatively well-studied GH29 and GH33 families are almost certain to function as fucosidases and sialidases, respectively, however, this level of confidence does not extend to less-explored families or those with diverse activities. Our analysis revealed 58 CAZyme (sub)families shared between all groups, with the GH13 family displaying the highest overall abundance (Fig 4b, Supplementary Table 8). The GH13 family is relatively well studied with substrate specificities and structures solved in many of the subfamilies. Notably for this study, this family solely has activity against α-linked glucose polysaccharides. The most prevalent of these polymers is plant-derived starch, which is likely why the GH13 family is so abundant, but others include glycogen (animal-derived), dextran (bacteria-derived) and pullulan (fungal-derived).

The *Bifidobacterium* genomes from Primates exhibited the greatest CAZyme diversity, encoding a total of 166 classes encompassing 34 unique families such as GH59, GH84, GH88, GH123, GH151 and CE12 (Fig 4b, Supplementary Table 8). Interestingly, all these unique GH families are associated with the breakdown of host glycans, including glycosphingolipids, O-glycans and glycosaminoglycans.

Conversely, *Bifidobacterium* genomes from Hymenoptera were the least diverse, with 87 classes and only 2 unique (sub)families - GH43_31 and GH144, which are associated with β-galactofuranosidase and β-glucanase activities, respectively. When analysing the abundance of specific CAZyme families, we observed distinct differences between host groups (Fig 4c). Primate-associated *Bifidobacterium* encoded the highest average number of GH enzymes and glycosyl transferases (GTs) (61.2 ± 20.7 and 23.7 ± 5.14 (mean ± sd), respectively), while the lowest abundances of these enzyme classes were recorded in *Bifidobacterium* from Artiodactyla (36.7 ± 9.08 GH enzymes) and Hymenoptera (18.1 ± 2.86 GTs), respectively. Notably, carbohydrate binding module (CBM) families were the most abundant in *Bifidobacterium* genomes associated with Artiodactyla (9.05 ± 2.56), while Rodentia and Carnivora showed similar numbers for carbohydrate esterases (CEs) (6.62 ± 1.87 and 6.45 ± 1.44, respectively).

To further assess the diversity of bifidobacterial CAZymes across host groups, we performed PCA on the CAZyme abundance matrix. The PCA did not reveal clear separation between different host groups (Fig 4d), which was supported by ANOSIM statistics (GH: R = 0.2975 p = 0.001 and ANOSIM CBM, CE, GT R: 0.3521, p = 0.001). However, additional analysis of the 189 CAZyme families and subfamilies in *Bifidobacterium* from Hymenoptera (insect-associated) vs. those from mammals, Primates, and non-primates (QBonferroni<0.05; Fig. 4e, Supplementary Table 9) indicated significant differences in CAZyme abundances. Compared to insects, the GH13 family was more prevalent in mammals. Within the GH13 family in mammals, subfamilies GH13_3 (glucan biosynthesis in bacteria), GH13_9 (glycogen branch synthesis), and GH13_11 (glycogen metabolism), were widely represented, with GH13_4 (amylosucrases) and GH13_20 significantly more abundant in Primates.

A representation of the GH family abundance matrix, normalised to the number of genomes per host group, revealed distinct patterns in the distribution of particular GH families across host groups, consistent with our association analysis (Supplementary Figure 1). The heatmap indicates a consistent difference between the putative CAZy GH families present in the insect-associated bifidobacterial genomes compared to the other four groups. For example, GH5_18 (β-mannose), GH29 (α-fucose), GH32 (levanases), GH38 (α-mannosidases), GH43_22 (α-arabinofuranosidases and β-xylosidases), GH65 (α-glucose), GH78 (α-rhamnosidase), GH146 (β-arabinofuranosidases) families are relatively more abundant in insects. Interestingly, many of these family are associated with breaking down plant glycans and polysaccharides. These findings highlight the functional adaptations of *Bifidobacterium* to different host environments, particularly in relation to carbohydrate metabolism.

### Structural and functional diversity of GH13 enzymes in *Bifidobacterium* across diverse animal hosts

GH13 family sequences were the most abundantly represented in our dataset and showed associations with specific animal host groups. Due to these unique attributes, we postulated that this GH13 enzymes could be used as a sentinel family for assessing evolutionary functional processes. We utilised 4303 amino acid sequences to construct a maximum-likelihood evolutionary tree (Supplementary Figure 2), which revealed well-defined clusters within GH13 subfamilies. The sequence diversity within these subfamilies was evident, as reflected by the presence of distinct subclusters, consistent with previous reports (66).

To understand how this sequence diversity translates into functional potential, we compared the predicted structures of representative proteins from GH13 subfamilies that were particularly abundant in bifidobacteria associated with mammals in comparison to insects. Specifically, we analysed the α-maltosyltransferase GH13_3, the α-glucan branching GH13_9 and the α-glucan debranching GH13_11 structures, revealing obvious differences in the presence of non-catalytic domains among enzymes from Primate-, Rodentia- and Artiodactyla-associated species (Fig. 5a). Superimposition of these structures with reference models complexed with their respective identified ligands (4U3C with maltohexaose, 5GQX with maltoheptaose, and 7U3B with acarbose, respectively), indicated that bifidobacterial enzymes likely have similar substrate affinities, with conserved ligand orientation and positioning across subfamilies – likely due to their function in glycogen metabolism.

**Fig 5.**
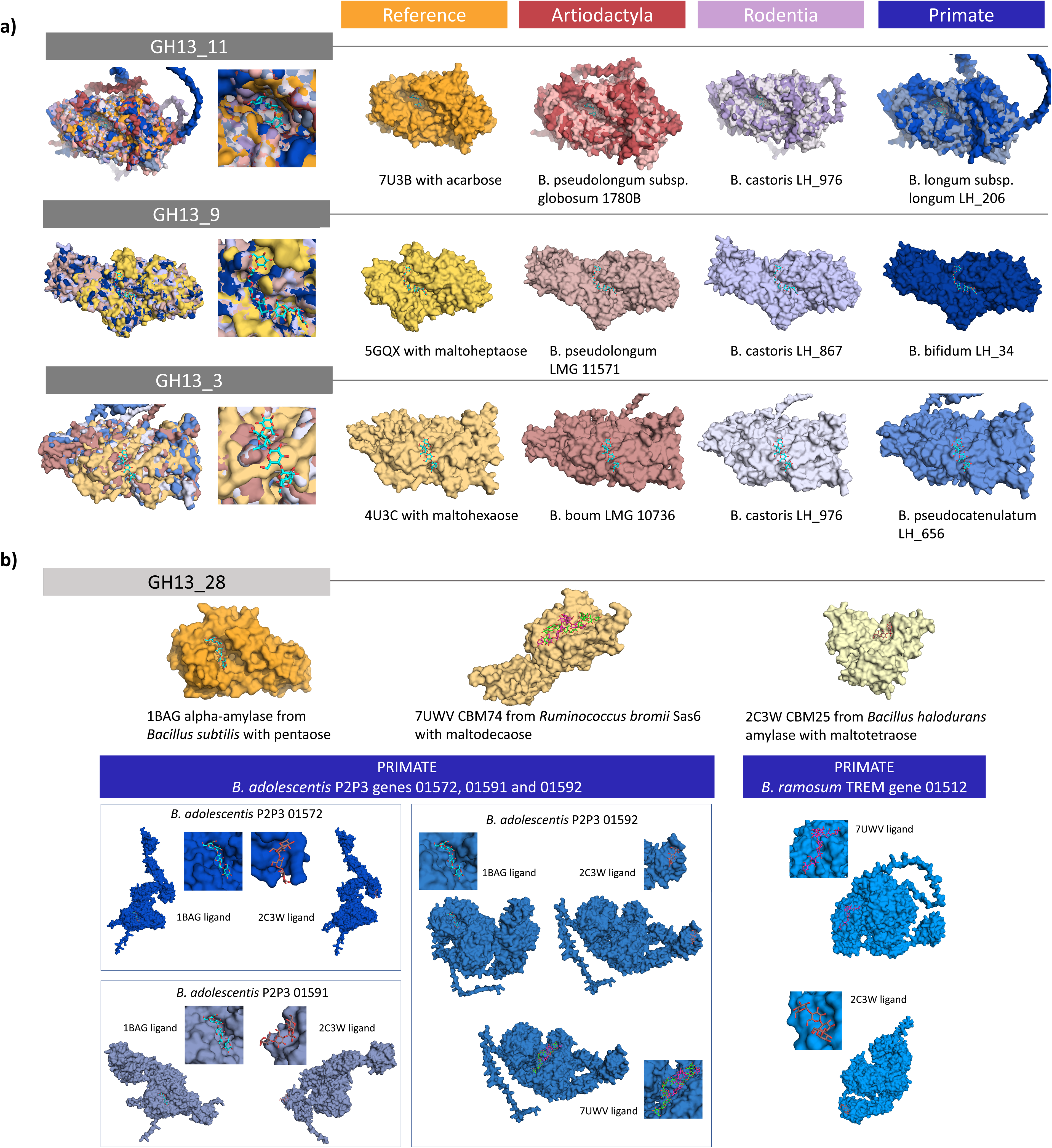
Analysis of bifidobacterial GH13 glycoside hydrolases. **a)** Comparison of AlphaFold models generated for selected bifidobacterial sequences with structures of representative proteins from subfamilies indicated as particularly abundant in bifidobacteria associated with mammals: GH13_11 (glycogen metabolism), GH13_9 and GH13_3 (glycogen synthesis). **b)** Comparison of AlphaFold models generated for GH13_28 sequences from selected primate bifidobacterial strains linked to the degradation of resistant starches – **Bifidobacterium* adolescentis* P2P3 and **Bifidobacterium* ramosum* TREM – with solved structures of selected GH13_28 proteins (1BAG), and specifically, starch-associated carbohydrate binding modules CBM25 (2C3W) and CBM74 (7UWV) coupled with their respective ligands. Protein models from primate-associated strains are coloured in the shades of blue, those from bifidobacteria isolated from hosts belonging to order Artiodactyla in the shades of red and those from rodent-associated **Bifidobacterium** in the shades of purple.

We also compared the structures of GH13 subfamilies significantly associated with Primates vs other mammals, including the GH13_4 amylosucrases and the GH13_20 (variety of activities), with a range of reference models complexed with their ligands (Supplementary Figure 3). Our analysis suggested that bifidobacterial enzymes from these subfamilies might bind multiple substrates. For instance, the pocket-shaped active site of GH13_4 amylosucrases could accommodate both sucrose and maltohexaose, while the open cleft-shaped active sites of GH13_20 enzymes likely prefer cyclodextrins, but may also bind other ligands, such as maltohexaose (B. psychraerophilum DSM 22366), and short-chain oligosaccharides (B. longum subsp. longum LH_12).

Notably, concerning the breakdown of α-linked glucose polysaccharides, mammalian-associated bifidobacteria are recognised as primary degraders of resistant starch (RS), with species such as *Bifidobacterium* adolescentis, *Bifidobacterium* choerinum and B. pseudolongum exhibiting RS-degrading phenotypes (67–69). RS has a compact crystalline structure, so the enzymes and binding modules that act on it have binding sites with very different conformations to those that have specificity for soluble substrates. This capacity is linked to GH13_28 α-amylases containing specific carbohydrate-binding modules (CBMs), namely CBM25, CBM26, and CBM74 (69). We identified GH13_28 enzymes containing CBM74 and either CBM25 and/or the CBM26 modules in 57 out of the 387 genomes (14.7%), spanning 20 bifidobacterial species and subspecies. Most of these genomes (84.2%, n=48) also contained additional α-amylases with either CBM25 or CBM26 modules alone. Further analysis revealed that some species, such as *Bifidobacterium* castoris, consistently possessed genes encoding GH13_28 α-amylases, while in other species, their presence was strain dependent.

We overlaid the structures of identified GH13_28 proteins from *Bifidobacterium* strains with reported RS-degrading phenotypes (B. adolescentis P2P3, B. choerinum FMB -1, and an in-house strain B. castoris LH_775) onto selected models of α-amylases (1BAG) and starch-recognising modules CBM25 (2C3W) and CBM74 (7UWV) complexed with α-glucans (Fig. 5b and Supplementary Figure 4). Successful overlay of different ligands onto the same structure allowed for the identification of the presence of both the catalytic domain and the carbohydrate binding module(s) in several of the selected bifidobacterial GH13_28 enzymes, confirming the presence of multiple carbohydrate-binding domains with specific substrate preferences. Binding sites for the 2C3W CBM25 ligand from Bacillus halodurans (maltotetraose) were conserved across all *Bifidobacterium* GH13_28 structures. Similarly, we identified binding sites for either double or single helical starch (model 7UWV from Ruminococcus bromii) in bifidobacterial structures containing a predicted CBM74 domain. These results were replicated for other animal-associated species identified as potential RS degraders based on genomic predictions, namely *Bifidobacterium* ramosum, *Bifidobacterium* thermophilum and *Bifidobacterium* tsurumiense (Fig. 5b and Supplementary Figure 4). Our findings suggest that CAZy annotations coupled with structural modelling can provide valuable insights into the RS-degrading phenotype of bifidobacterial strains.

## Discussion

By combining taxonomic profiling, genomic, phylogenetic, and functional analyses, we demonstrate how host phylogeny and ecology shape the composition, function, and evolutionary trajectories of *Bifidobacterium* at a strain-resolved level.

Our data indicates that gut microbiome composition is significantly influenced by host phylogeny, particularly in mammals. At the strain level, *Bifidobacterium* displayed strong host-specific co- phylogenetic associations, likely driven by vertical transmission and dietary factors. These results align with previous studies which have suggested strong signals of phylosymbiosis between mammalian hosts and their gut microbiota (70,71), yet our dataset extends these findings by showing that associations of *Bifidobacterium* are stronger in hosts with specific dietary niches. For instance, high *Bifidobacterium* abundance and strong phylogenetic signals in Primates, particularly marmosets and tamarins, are likely driven by diets rich in complex carbohydrates and fibrous plant material (72), resulting in persistence of bifidobacterial species adept at glycan utilisation. Additionally, the rise of Veillonellales-Selenomonadales in the ancestors of the New World monkeys, positively covarying with Bifidobacteriales, suggests potential cross-feeding interactions. This mirrors interactions observed in humans, where scavenging Veillonella species ferment lactate produced by B. longum subsp. infantis to acetate and propionate, indicating conserved microbial interaction networks across hosts (73).

A key finding was the robust co-phylogenetic signal between rodents and their *Bifidobacterium* strains, despite our earlier studies suggesting no such relationship in wild mice (genus Apodemus) (74). This shift in observations likely stems from the expanded dataset and different methodological approach in our current study, which included a host timetree, a broader range of *Bifidobacterium* strains and the GTDB-tk bac120 marker set, and highlights the importance of comprehensive genomic datasets and robust phylogenetic methods in revealing patterns of host-microbe co- evolution (75). These results also underscore the importance of vertical transmission in maintaining phylogenetic congruence at the strain level and suggest that co-evolution between *Bifidobacterium* and their hosts is nuanced and complex. The evolutionary stability of these host-microbe relationships at the strain level suggests that vertical transmission potentially overshadows ecological factors such as cross-species transmission, and raises important questions about the mechanisms driving phylosymbiosis, including the potential role of additional factors such as microbial interactions with the host immune system or strain retention due to nutrient availability.

In contrast, we observed a weaker phylosymbiosis signal in birds, likely reflecting the more variable nature of their gut microbiota (76). Although the smaller sample size limits our conclusions here, the ancestral reconstruction of microbiota shifts in bird lineages, particularly the increased abundance of Enterobacteriales and Clostridiales in Neoaves, mirrors the dietary shifts towards carnivory in Carnivora. These dietary changes appear to be a driving force behind the microbiota shifts we observed in both avian and mammalian lineages (77). Interestingly, a correlation between species of Clostridium and animal-based diet has also been observed in humans (78). Members of this genus have previously been identified as important degraders of amino acids, including lysine, alanine and glycine, particularly abundant in meat and meat products (79,80). Overall, our findings point to a complex interplay between host phylogeny, diet, and vertical transmission, rather than purely ecological dynamics driving these patterns.

Our functional analyses revealed striking host-specific metabolic adaptations in *Bifidobacterium*, particularly in carbohydrate metabolism and oxidative stress responses. Insect-associated *Bifidobacterium* were enriched in oxidative phosphorylation genes. This finding is particularly intriguing given the anoxic nature of insect guts (81), suggesting that bifidobacterial cytochrome bd oxidases might play a yet unclear role. One possibility is defence against reactive oxygen species (ROS) released into the insect gut environment as part of the innate immune system; a mechanism which has also been seen in E. coli and Mycobacterium smegmatis (82,83). These findings raise the possibility that *Bifidobacterium* has evolved unique mechanisms to mitigate oxidative stress, an adaptation that could represent a previously unappreciated aspect of insect gut symbiosis.

In mammals, the diversity and abundance of bifidobacterial GH enzymes, particularly those involved in glycogen metabolism (GH13_9 and GH13_11), reflect adaptations to complex carbohydrate-rich diets. Consistent with previous findings, our study also shows the absence of these functions in insect-associated *Bifidobacterium*, likely reflecting the nectar and pollen-based diets of their hosts, which lack complex α-glucoside linkages (84,85). The absence of glycogen metabolism suggests an evolutionary loss, potentially due to reductive genome evolution; a phenomenon previously observed in obligate intracellular symbionts (86,87). However, since insect-associated *Bifidobacterium* are not obligate symbionts, these strains may rely on glycogen metabolism by the host or other microbiota members, highlighting an example of microbial interdependence within the gut ecosystem.

Moreover, the identification of the presence of multi-domain GH13_28 α-amylases containing CBM25, CBM26, and the relatively novel CBM74 module in *Bifidobacterium* across various hosts highlights the genus’s ecological flexibility. As recently demonstrated in Ruminococcus bromii (88), enzymes containing these modules are crucial for the efficient degradation of both short- and long-chain starch molecules, including RS, which are prevalent in herbivorous diets. Indeed, our structural modelling confirms the affinity of these carbohydrate-binding domains for their respective ligands, highlighting the metabolic versatility of *Bifidobacterium*. This underscores the adaptive potential of *Bifidobacterium* to diverse dietary environments and suggests potential targets for enhancing their starch-degrading capabilities in animal and human probiotic applications.

In conclusion, this research significantly broadens our understanding of the evolutionary biology and functional ecology of *Bifidobacterium*. By integrating diverse approaches, we have revealed how the wider microbiome and these specific microbes adapt to their hosts through a balance of vertical transmission, dietary influences, and host-specific interactions. This work provides a foundation for targeted therapeutic interventions tailored to specific host diets or ecological niches. Future research should explore how dietary and ecological components influence microbiota across hosts, enhancing our understanding of gut microbiota dynamics and their functional roles in health and disease.

## Data availability

16S rRNA amplicon sequencing data analysed in this study have been deposited to the NCBI SRA under the SRA BioProject number PRJNA1200941. The draft genomes of 96 *Bifidobacterium* isolates sequenced here have been deposited to the NCBI Genome database under the BioProject number PRJNA1200594.

## Acknowledgments

We would like to thank the zoo staff at Banham Zoo (UK), Africa Alive! (UK) and Pafos Zoo (Cyprus) for their assistance in collecting samples for this study.

## Author contributions

LJH and MK designed the overall study. SG and IC were responsible for sample collection. JK and LC processed faecal samples and extracted DNA for the 16S rRNA amplicon sequencing. MK pre-processed the 16S rRNA sequencing reads. SM performed initial analyses of the 16S rRNA sequencing data. Taxonomic analyses were performed by DS and phylosymbiosis analysis was performed by MK. JK and MK processed faecal samples for bacterial isolations, isolated *Bifidobacterium* strains and extracted genomic DNA for WGS. MK performed all genomic, evolutionary and functional prediction analysis and visualised the data. MK and LJH drafted the manuscript, DS, and LC provided further edits and co-writing of the final version. All authors read and approved the final manuscript.

## References

1. Gotoh A, Katoh T, Sakanaka M, Ling YW, Yamada C, Asakuma S, et al. Sharing of human milk oligosaccharides degradants within bifidobacterial communities in faecal cultures supplemented with *Bifidobacterium* bifidum. Sci Rep. 2018 Sep 18;8:13958.

2. Thomson P, Medina DA, Garrido D. Human milk oligosaccharides and infant gut bifidobacteria: Molecular strategies for their utilization. Food Microbiol. 2018 Oct;75:37–46.

3. Lawson MAE, O’Neill IJ, Kujawska M, Gowrinadh Javvadi S, Wijeyesekera A, Flegg Z, et al. Breast milk-derived human milk oligosaccharides promote *Bifidobacterium* interactions within a single ecosystem. Isme J. 2020 Feb;14(2):635–48.

4. Milani C, Turroni F, Duranti S, Lugli GA, Mancabelli L, Ferrario C, et al. Genomics of the Genus *Bifidobacterium* Reveals Species-Specific Adaptation to the Glycan-Rich Gut Environment. Appl Environ Microbiol. 2016 Feb;82(4):980–91.

5. Lugli GA, Milani C, Turroni F, Duranti S, Mancabelli L, Mangifesta M, et al. Comparative genomic and phylogenomic analyses of the Bifidobacteriaceae family. BMC Genomics. 2017 Aug 1;18.

6. Lugli GA, Mancino W, Milani C, Duranti S, Mancabelli L, Napoli S, et al. Dissecting the Evolutionary Development of the Species *Bifidobacterium* animalis through Comparative Genomics Analyses. Appl Environ Microbiol. 2019 Apr 1;85(7):e02806–18.

7. Lugli GA, Duranti S, Albert K, Mancabelli L, Napoli S, Viappiani A, et al. Unveiling Genomic Diversity among Members of the Species *Bifidobacterium* pseudolongum, a Widely Distributed Gut Commensal of the Animal Kingdom. Appl Environ Microbiol. 2019 Apr;85(8):e03065–18.

8. Milani C, Mangifesta M, Mancabelli L, Lugli GA, James K, Duranti S, et al. Unveiling bifidobacterial biogeography across the mammalian branch of the tree of life. Isme J. 2017 Dec;11(12):2834–47.

9. Sayers EW, Beck J, Brister JR, Bolton EE, Canese K, Comeau DC, et al. Database resources of the National Center for Biotechnology Information. Nucleic Acids Res. 2020 Jan 8;48(D1):D9–16.

10. Orkin JD, Campos FA, Myers MS, Cheves Hernandez SE, Guadamuz A, Melin AD. Seasonality of the gut microbiota of free-ranging white-faced capuchins in a tropical dry forest. ISME J. 2019 Jan;13(1):183–96.

11. Modrackova N, Stovicek A, Burtscher J, Bolechova P, Killer J, Domig KJ, et al. The bifidobacterial distribution in the microbiome of captive primates reflects parvorder and feed specialization of the host. Sci Rep. 2021 Jul 27;11(1):15273.

12. Malukiewicz J, D’arc M, Dias CA, Cartwright RA, Grativol AD, Moreira SB, et al. Bifidobacteria define gut microbiome profiles of golden lion tamarin (Leontopithecus rosalia) and marmoset (Callithrix sp.) metagenomic shotgun pools. Sci Rep. 2023 Sep 21;13(1):15679.

13. Lim SJ, Bordenstein SR. An introduction to phylosymbiosis. Proc Biol Sci. 2020 Mar 11;287(1922):20192900.

14. Legendre P, Desdevises Y, Bazin E. A statistical test for host-parasite coevolution. Syst Biol. 2002 Apr;51(2):217–34.

15. Page RDM. Tangled Trees: Phylogeny, Cospeciation, and Coevolution. University of Chicago Press; 2003. 361 p.

16. Balbuena JA, Míguez-Lozano R, Blasco-Costa I. PACo: A Novel Procrustes Application to Cophylogenetic Analysis. PLOS ONE. 2013 Apr 8;8(4):e61048.

17. Moeller AH, Caro-Quintero A, Mjungu D, Georgiev AV, Lonsdorf EV, Muller MN, et al. Cospeciation of gut microbiota with hominids. Science. 2016 Jul 22;353(6297):380–2.

18. Lugli GA, Alessandri G, Milani C, Mancabelli L, Ruiz L, Fontana F, et al. Evolutionary development and co-phylogeny of primate-associated bifidobacteria. Environ Microbiol. 2020 Aug;22(8):3375–93.

19. Lugli GA, Milani C, Duranti S, Mancabelli L, Mangifesta M, Turroni F, et al. Tracking the taxonomy of the genus *Bifidobacterium* based on a phylogenomic approach. Appl Environ Microbiol. 2017 Dec 8;84(4):e02249–17.

20. Rodriguez CI, Martiny JBH. Evolutionary relationships among bifidobacteria and their hosts and environments. BMC Genomics. 2020 Jan 8;21(1):26.

21. Berkhout MD, Plugge CM, Belzer C. How microbial glycosyl hydrolase activity in the gut mucosa initiates microbial cross-feeding. Glycobiology. 2021 Oct 18;32(3):182–200.

22. La Rosa SL, Ostrowski MP, Vera-Ponce de León A, McKee LS, Larsbrink J, Eijsink VG, et al. Glycan processing in gut microbiomes. Curr Opin Microbiol. 2022 Jun 1;67:102143.

23. Alessandri G, van Sinderen D, Ventura M. The genus *Bifidobacterium*: From genomics to functionality of an important component of the mammalian gut microbiota running title: Bifidobacterial adaptation to and interaction with the host. Comput Struct Biotechnol J. 2021 Mar 9;19:1472–87.

24. Walker AW, Martin JC, Scott P, Parkhill J, Flint HJ, Scott KP. 16S rRNA gene-based profiling of the human infant gut microbiota is strongly influenced by sample processing and PCR primer choice. Microbiome. 2015 Jun 22;3:26.

25. Suzuki MT, Giovannoni SJ. Bias caused by template annealing in the amplification of mixtures of 16S rRNA genes by PCR. Appl Environ Microbiol. 1996 Feb;62(2):625.

26. Kozich JJ, Westcott SL, Baxter NT, Highlander SK, Schloss PD. Development of a Dual-Index Sequencing Strategy and Curation Pipeline for Analyzing Amplicon Sequence Data on the MiSeq Illumina Sequencing Platform. Appl Environ Microbiol. 2013 Sep;79(17):5112.

27. Martin M. Cutadapt removes adapter sequences from high-throughput sequencing reads. EMBnet.journal. 2011 May 2;17(1):10–2.

28. Callahan BJ, McMurdie PJ, Rosen MJ, Han AW, Johnson AJA, Holmes SP. DADA2: High-resolution sample inference from Illumina amplicon data. Nat Methods. 2016 Jul;13(7):581–3.

29. Quast C, Pruesse E, Yilmaz P, Gerken J, Schweer T, Yarza P, et al. The SILVA ribosomal RNA gene database project: improved data processing and web-based tools. Nucleic Acids Res. 2013 Jan 1;41(D1):D590–6.

30. Andersen KS, Kirkegaard RH, Karst SM, Albertsen M. ampvis2: an R package to analyse and visualise 16S rRNA amplicon data [Internet]. bioRxiv; 2018 [cited 2024 Jun 6]. p. 299537. Available from: https://www.biorxiv.org/content/10.1101/299537v1

31. Team RC. R: A language and environment for statistical computing. R Found Stat Comput. 2017;Available at: http://www.r-project.org/index.html.

32. Oksanen J, Blanchet FG, Friendly M, Kindt r, Legendre P, McGillin D, et al. vegan: Community Ecology Package. R Package Version 25–6. 2019;https://CRAN.R-project.org/package=vegan.

33. Kolde R. pheatmap: Pretty Heatmaps [Internet]. 2019 [cited 2024 Nov 14]. Available from: https://cran.r-project.org/web/packages/pheatmap/index.html

34. Wickham H, Chang W, Henry L, Pedersen TL, Takahashi K, Wilke C, et al. ggplot2: Create Elegant Data Visualisations Using the Grammar of Graphics [Internet]. 2024 [cited 2024 Jun 10]. Available from: https://cran.r-project.org/web/packages/ggplot2/index.html

35. Kumar S, Stecher G, Suleski M, Hedges SB. TimeTree: A Resource for Timelines, Timetrees, and Divergence Times. Mol Biol Evol. 2017 Jul 1;34(7):1812–9.

36. Perez-Lamarque B, Sommeria-Klein G, Duret L, Morlon H. Phylogenetic Comparative Approach Reveals Evolutionary Conservatism, Ancestral Composition, and Integration of Vertebrate Gut Microbiota. Mol Biol Evol. 2023 Jul 1;40(7):msad144.

37. Guo J, Gabry J, Goodrich B, Johnson A, Weber S, Badr HS, et al. rstan: R Interface to Stan [Internet]. 2024 [cited 2024 Jun 10]. Available from: https://cran.r-project.org/web/packages/rstan/index.html

38. Page AJ, De Silva N, Hunt M, Quail MA, Parkhill J, Harris SR, et al. Robust high-throughput prokaryote de novo assembly and improvement pipeline for Illumina data. Microb Genomics. 2016 Aug 25;2(8):e000083.

39. Chen S, Zhou Y, Chen Y, Gu J. fastp: an ultra-fast all-in-one FASTQ preprocessor. Bioinformatics. 2018 Sep 1;34(17):i884–90.

40. Wick RR, Judd LM, Gorrie CL, Holt KE. Unicycler: Resolving bacterial genome assemblies from short and long sequencing reads. PLoS Comput Biol. 2017 Jun;13(6):e1005595.

41. Parks DH, Imelfort M, Skennerton CT, Hugenholtz P, Tyson GW. CheckM: assessing the quality of microbial genomes recovered from isolates, single cells, and metagenomes. Genome Res. 2015;25(7):1043–55.

42. Olm MR, Brown CT, Brooks B, Banfield JF. dRep: a tool for fast and accurate genomic comparisons that enables improved genome recovery from metagenomes through de-replication. ISME J. 2017 Dec 1;11(12):2864–8.

43. Chaumeil PA, Mussig AJ, Hugenholtz P, Parks DH. GTDB-Tk v2: memory friendly classification with the genome taxonomy database. Bioinformatics. 2022 Nov 30;38(23):5315–6.

44. Pritchard L, Glover RH, Humphris S, Elphinstone JG, Toth IK. Genomics and taxonomy in diagnostics for food security: soft-rotting enterobacterial plant pathogens. Anal Methods. 2016;8(1):12–24.

45. Chun J, Oren A, Ventosa A, Christensen H, Arahal DR, da Costa MS, et al. Proposed minimal standards for the use of genome data for the taxonomy of prokaryotes. Int J Syst Evol Microbiol. 2018;68(1):461–6.

46. Minh BQ, Schmidt HA, Chernomor O, Schrempf D, Woodhams MD, von Haeseler A, et al. IQ-TREE 2: New Models and Efficient Methods for Phylogenetic Inference in the Genomic Era. Mol Biol Evol. 2020 May 1;37(5):1530–4.

47. Letunic I, Bork P. Interactive Tree of Life (iTOL) v6: recent updates to the phylogenetic tree display and annotation tool. Nucleic Acids Res. 2024 Apr 13;gkae268.

48. Paradis E, Schliep K. ape 5.0: an environment for modern phylogenetics and evolutionary analyses in R. Bioinformatics. 2019 Feb 1;35(3):526–8.

49. Revell LJ. phytools: an R package for phylogenetic comparative biology (and other things). Methods Ecol Evol. 2012;3(2):217–23.

50. Seemann T. Prokka: rapid prokaryotic genome annotation. Bioinformatics. 2014 Jul 15;30(14):2068–9.

51. Huerta-Cepas J, Forslund K, Coelho LP, Szklarczyk D, Jensen LJ, von Mering C, et al. Fast Genome-Wide Functional Annotation through Orthology Assignment by eggNOG-Mapper. Mol Biol Evol. 2017 Aug;34(8):2115–22.

52. Huerta-Cepas J, Szklarczyk D, Heller D, Hernandez-Plaza A, Forslund SK, Cook H, et al. eggNOG 5.0: a hierarchical, functionally and phylogenetically annotated orthology resource based on 5090 organisms and 2502 viruses. Nucleic Acids Res. 2019 Jan 8;47(D1):D309–14.

53. Wickham H, François R, Henry L, Müller K, Vaughan D, Software P, et al. dplyr: A Grammar of Data Manipulation [Internet]. 2023 [cited 2024 Jun 10]. Available from: https://cran.r-project.org/web/packages/dplyr/index.html

54. Rühlemann MC, Bang C, Gogarten JF, Hermes BM, Groussin M, Waschina S, et al. Functional host-specific adaptation of the intestinal microbiome in hominids. Nat Commun. 2024 Jan 6;15(1):326.

55. Wilke CO. cowplot: Streamlined Plot Theme and Plot Annotations for ‘ggplot2’ [Internet]. 2024 [cited 2024 Jun 10]. Available from: https://cran.r-project.org/web/packages/cowplot/index.html

56. Conway J, Gehlenborg N. UpSetR: A More Scalable Alternative to Venn and Euler Diagrams for Visualizing Intersecting Sets [Internet]. 2019 [cited 2024 Jun 10]. Available from: https://cran.r-project.org/web/packages/UpSetR/index.html

57. Kassambara A, Mundt F. factoextralll: Extract and Visualize the Results of Multivariate Data Analyses. 2020;Available at: http://www.sthda.com/english/rpkgs/factoextra.

58. Berman HM, Westbrook J, Feng Z, Gilliland G, Bhat TN, Weissig H, et al. The Protein Data Bank. Nucleic Acids Res. 2000 Jan 1;28(1):235–42.

59. Emsley P, Lohkamp B, Scott WG, Cowtan K. Features and development of Coot. Acta Crystallogr D Biol Crystallogr. 2010 Apr;66(Pt 4):486–501.

60. DeLano WL. The PyMOL Molecular Graphics System. Schrödinger, LLC;

61. Bibekar P, Krapp L, Peraro MD. PeSTo-Carbs: Geometric Deep Learning for Prediction of Protein–Carbohydrate Binding Interfaces. J Chem Theory Comput. 2024 Apr 11;20(8):2985–91.

62. Hird SM, Sánchez C, Carstens BC, Brumfield RT. Comparative gut microbiota of 59 neotropical bird species. Front Microbiol 6: 1403. 2015.

63. Amato KR, G. Sanders J, Song SJ, Nute M, Metcalf JL, Thompson LR, et al. Evolutionary trends in host physiology outweigh dietary niche in structuring primate gut microbiomes. Isme J. 2019 Mar;13(3):576–87.

64. Brooks AW, Kohl KD, Brucker RM, van Opstal EJ, Bordenstein SR. Phylosymbiosis: Relationships and Functional Effects of Microbial Communities across Host Evolutionary History. PLoS Biol. 2016 Nov 18;14(11):e2000225.

65. Kanehisa M, Sato Y, Kawashima M, Furumichi M, Tanabe M. KEGG as a reference resource for gene and protein annotation. Nucleic Acids Res. 2016 Jan 4;44(D1):D457–462.

66. Stam MR, Danchin EG, Rancurel C, Coutinho PM, Henrissat B. Dividing the large glycoside hydrolase family 13 into subfamilies: towards improved functional annotations of alpha-amylase-related proteins. Protein Eng Des Sel. 2006 Dec;19(12):555–62.

67. Jung DH, Seo DH, Kim GY, Nam YD, Song EJ, Yoon S, et al. The effect of resistant starch (RS) on the bovine rumen microflora and isolation of RS-degrading bacteria. Appl Microbiol Biotechnol. 2018 Jun 1;102(11):4927–36.

68. Jung DH, Kim GY, Kim IY, Seo DH, Nam YD, Kang H, et al. *Bifidobacterium* adolescentis P2P3, a Human Gut Bacterium Having Strong Non-Gelatinized Resistant Starch-Degrading Activity. 2019 Dec 28;29(12):1904–15.

69. Jung DH, Park CS. Resistant starch utilization by *Bifidobacterium*, the beneficial human gut bacteria. Food Sci Biotechnol. 2023 Jan 27;32(4):441–52.

70. Groussin M, Mazel F, Sanders JG, Smillie CS, Lavergne S, Thuiller W, et al. Unraveling the processes shaping mammalian gut microbiomes over evolutionary time. Nat Commun. 2017 Feb 23;8:14319.

71. Nishida AH, Ochman H. Rates of Gut Microbiome Divergence in Mammals. Mol Ecol. 2018 Apr;27(8):1884–97.

72. Susanne R, Ann-Kathrin O. Husbandry and Management of New World Species: Marmosets and Tamarins. Lab Primate. 2007 Sep 2;145.

73. Button JE, Cosetta CM, Reens AL, Brooker SL, Rowan-Nash AD, Lavin RC, et al. Precision modulation of dysbiotic adult microbiomes with a human-milk-derived synbiotic reshapes gut microbial composition and metabolites. Cell Host Microbe. 2023 Sep 13;31(9):1523–1538.e10.

74. Kujawska M, Raulo A, Millar M, Warren F, Baltrūnaitė L, Knowles SCL, et al. *Bifidobacterium* castoris strains isolated from wild mice show evidence of frequent host switching and diverse carbohydrate metabolism potential. ISME Commun. 2022 Feb 25;2(1):1–14.

75. Hayward A, Poulin R, Nakagawa S. A broadscale analysis of host-symbiont cophylogeny reveals the drivers of phylogenetic congruence. Ecol Lett. 2021;24(8):1681–96.

76. Grond K, Sandercock BK, Jumpponen A, Zeglin LH. The avian gut microbiota: community, physiology and function in wild birds. J Avian Biol. 2018;49(11):e01788.

77. Youngblut ND, Reischer GH, Walters W, Schuster N, Walzer C, Stalder G, et al. Host diet and evolutionary history explain different aspects of gut microbiome diversity among vertebrate clades. Nat Commun. 2019 May 16;10(1):2200.

78. Asnicar F, Berry SE, Valdes AM, Nguyen LH, Piccinno G, Drew DA, et al. Microbiome connections with host metabolism and habitual diet from 1,098 deeply phenotyped individuals. Nat Med. 2021 Jan 11;27(2):321.

79. Smith EA, Macfarlane GT. Enumeration of amino acid fermenting bacteria in the human large intestine: effects of pH and starch on peptide metabolism and dissimilation of amino acids. FEMS Microbiol Ecol. 1998 Apr 1;25(4):355–68.

80. Dai Z, Zheng W, Locasale JW. Amino acid variability, tradeoffs and optimality in human diet. Nat Commun. 2022 Nov 5;13(1):6683.

81. Zheng H, Powell JE, Steele MI, Dietrich C, Moran NA. Honeybee gut microbiota promotes host weight gain via bacterial metabolism and hormonal signaling. Proc Natl Acad Sci U S A. 2017 May 2;114(18):4775–80.

82. Lindqvist A, Membrillo-Hernández J, Poole RK, Cook GM. Roles of respiratory oxidases in protecting Escherichia coli K12 from oxidative stress. Antonie Van Leeuwenhoek. 2000 Jul 1;78(1):23–31.

83. Lu P, Heineke MH, Koul A, Andries K, Cook GM, Lill H, et al. The cytochrome bd-type quinol oxidase is important for survival of Mycobacterium smegmatis under peroxide and antibiotic-induced stress. Sci Rep. 2015 May 27;5(1):10333.

84. Bottacini F, Milani C, Turroni F, Sánchez B, Foroni E, Duranti S, et al. *Bifidobacterium* asteroides PRL2011 Genome Analysis Reveals Clues for Colonization of the Insect Gut. PLoS ONE. 2012 Sep 20;7(9):e44229.

85. Sun Z, Zhang W, Guo C, Yang X, Liu W, Wu Y, et al. Comparative genomic analysis of 45 type strains of the genus *Bifidobacterium*: a snapshot of its genetic diversity and evolution. Plos One. 2015;10(2):e0117912.

86. Henrissat B, Deleury E, Coutinho PM. Glycogen metabolism loss: a common marker of parasitic behaviour in bacteria? Trends Genet. 2002 Sep 1;18(9):437–40.

87. Esteban-Torres M, Ruiz L, Rossini V, Nally K, van Sinderen D. Intracellular glycogen accumulation by human gut commensals as a niche adaptation trait. Gut Microbes. 2023 Dec 31;15(1):2235067.

88. Photenhauer AL, Villafuerte-Vega RC, Cerqueira FM, Armbruster KM, Mareček F, Chen T, et al. The Ruminococcus bromii amylosome protein Sas6 binds single and double helical α-glucan structures in starch. Nat Struct Mol Biol. 2024 Feb;31(2):255–65.

