## Supplementary Figure 1 for "Host-specific microbiome and genomic signatures in *Bifidobacterium* reveal co-evolutionary and functional adaptations across diverse animal hosts"

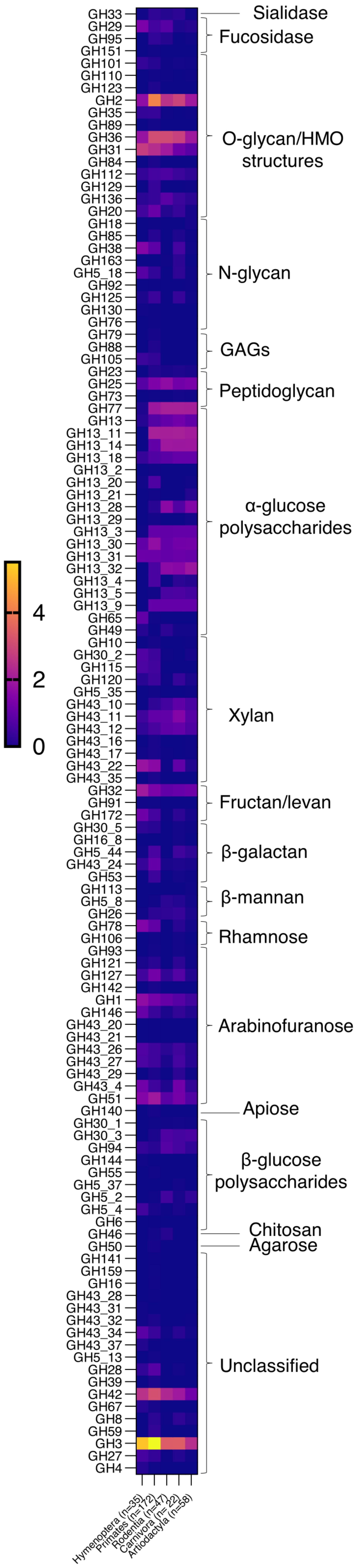


Supplementary Figure 1. Summarised representation of the glycoside hydrolase family abundance matrices obtained from dbCAN3 analysis. Results for individual *Bifidobacterium* genomes were grouped according to the host taxonomic level of order and normalised to the number of genomes per host group.
