## Supplementary Figure 2 for "Host-specific microbiome and genomic signatures in *Bifidobacterium* reveal co-evolutionary and functional adaptations across diverse animal hosts"

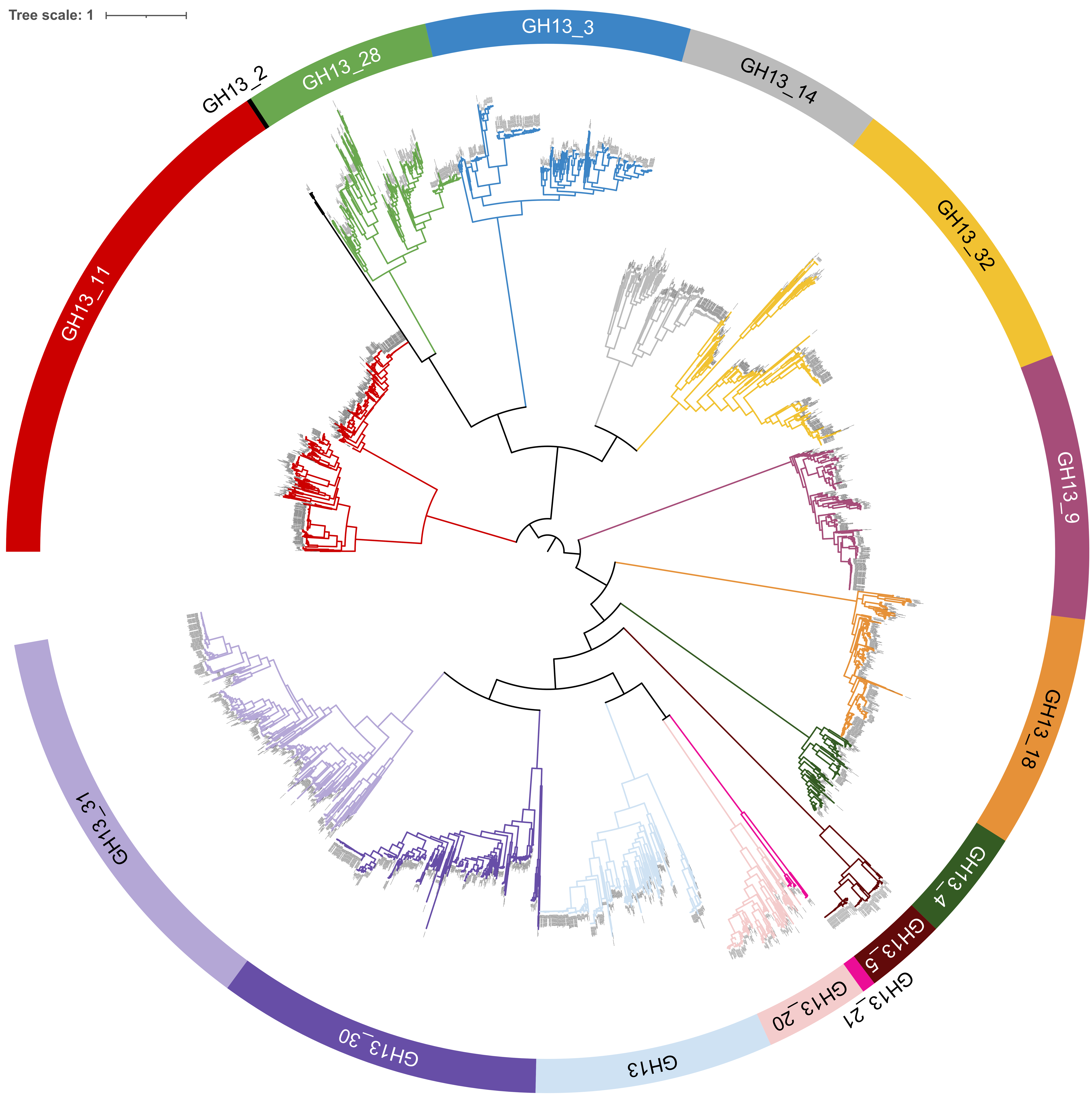


Supplementary Figure 2. Maximum likelihood phylogeny constructed based on 4303 bifidobacterial amino acid sequences identified in dbCAN3 analysis as belonging to the GH13 family of carbohydrate active enzymes (CAZymes) (amino acid substitution model WAG+F+R10). Different colours represent identified GH13 subfamilies.
