## Supplementary Figure 3 for "Host-specific microbiome and genomic signatures in *Bifidobacterium* reveal co-evolutionary and functional adaptations across diverse animal hosts"

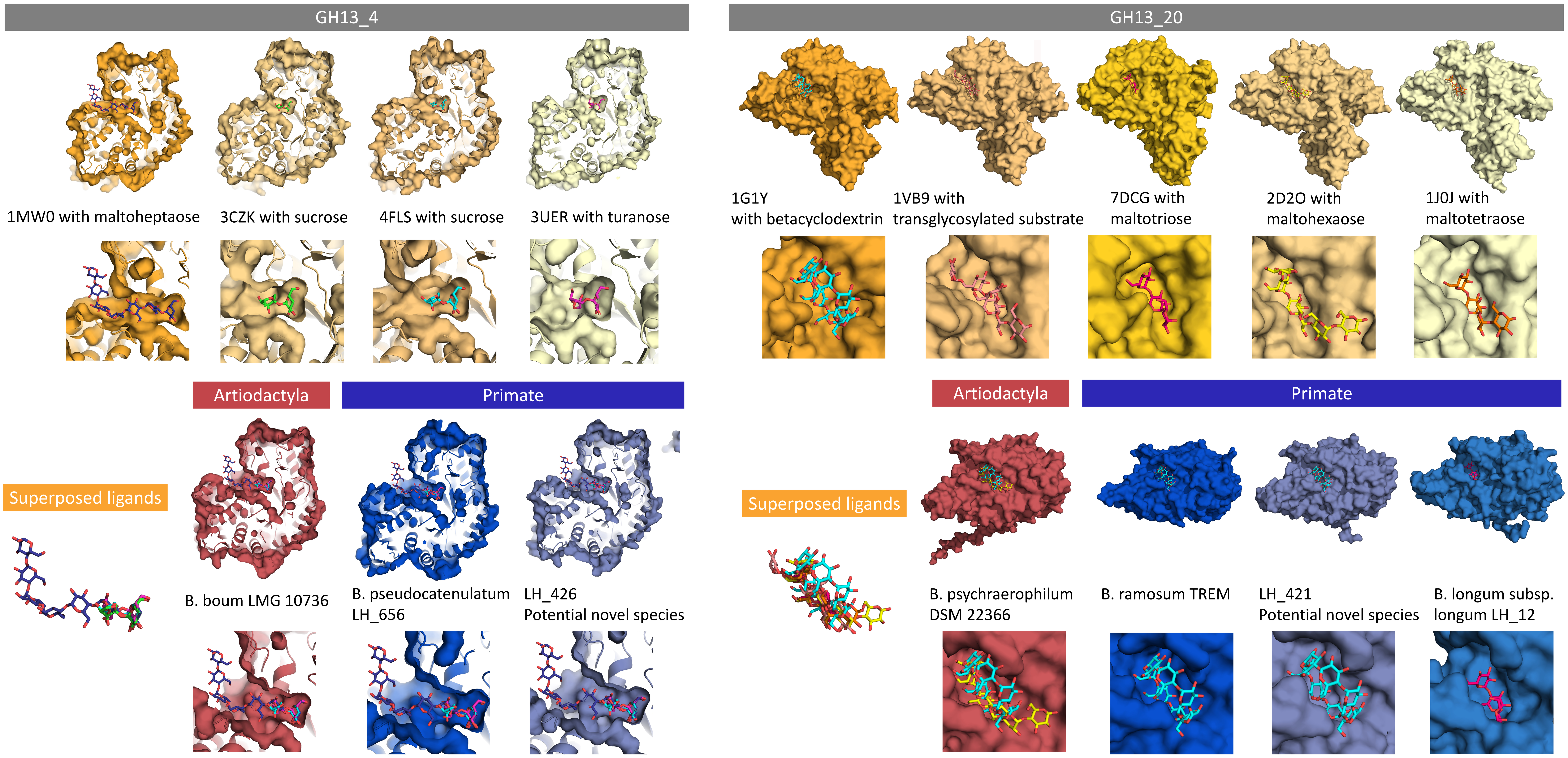


Supplementary Figure 3. Analysis of bifidobacterial GH13 glycoside hydrolases – comparison of AlphaFold models generated for selected bifidobacterial sequences belonging to subfamilies significantly associated with Primates – GH13_4 (extension of glycogen) and GH13_20 (glycogen metabolism) – with models of solved protein structures belonging to these GH subfamilies complexed with their respective ligands. Reference structures are represented in the shades of yellow. Protein models from primate-associated *Bifidobacterium* strains are coloured in the shades of blue and those from bifidobacteria isolated from hosts belonging to order Artiodactyla in the shades of red.
