## Supplementary Figure 4 for "Host-specific microbiome and genomic signatures in *Bifidobacterium* reveal co-evolutionary and functional adaptations across diverse animal hosts"

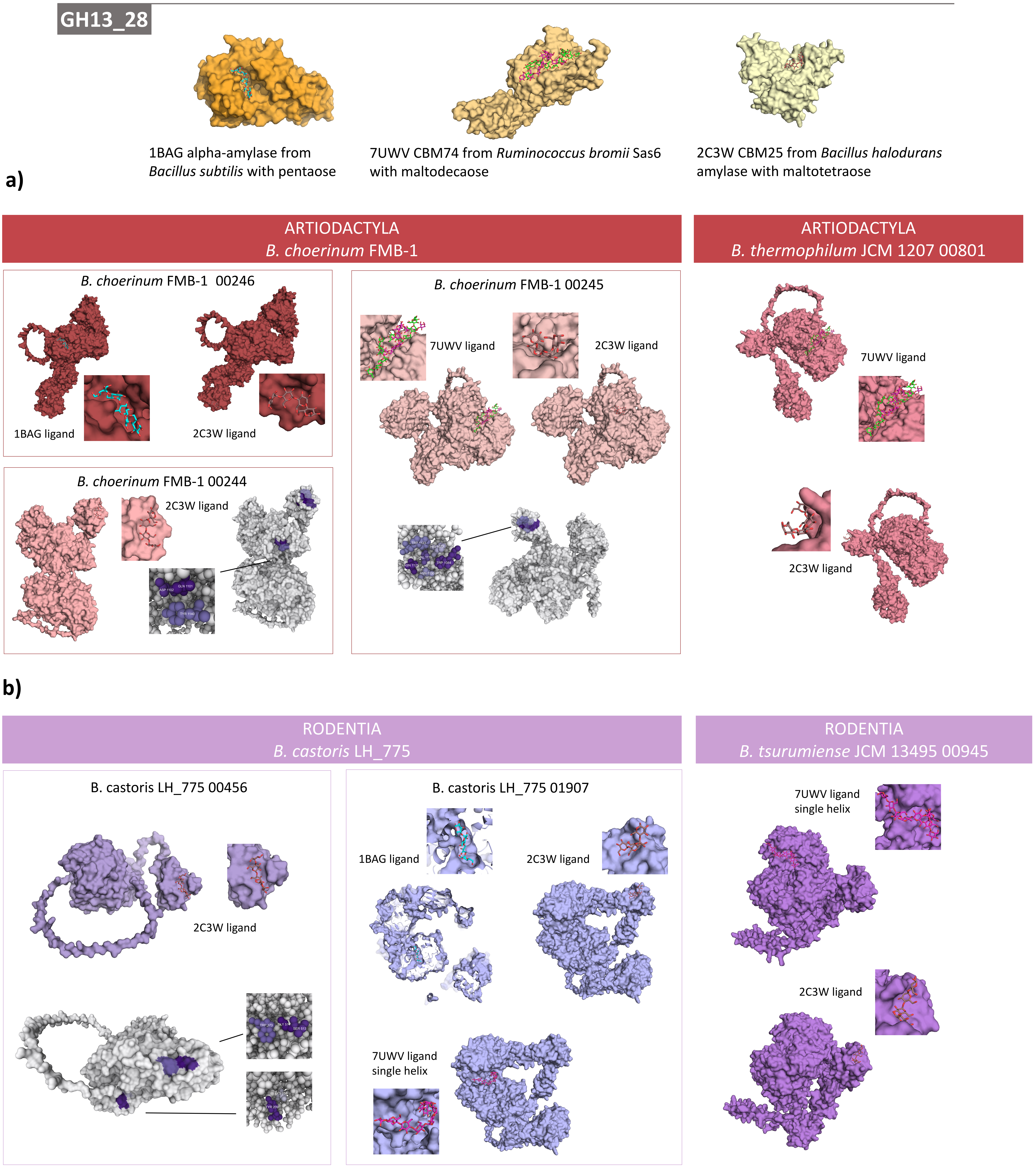


Supplementary Figure 4. Analysis of the bifidobacterial glycoside hydrolases belonging to the subfamily GH13_28 – comparison of AlphaFold models generated for GH13_28 sequences from selected bifidobacterial strains linked to the degradation of resistant starches with solved structures of selected GH13_28 proteins (1BAG), and specifically, starch-associated carbohydrate binding modules CBM25 (2C3W) and CBM74 (7UWV) coupled with their respective ligands. **a)** Representation of models from selected *Bifidobacterium* strains isolated from hosts belonging to order Artiodactyla (shades of red) and **b)** representation of models from selected rodent-associated *Bifidobacterium* (shades of purple). Grey representations show predicted carbohydrate binding sites, into which we were not able to fit ligands from used reference models.
